# Induced drought strongly affects richness and composition of ground-dwelling ants in the eastern Amazon

**DOI:** 10.1101/2020.08.10.242180

**Authors:** Rony P.S. Almeida, Rogério R. Silva, Antonio C.L. da Costa, Leandro V. Ferreira, Patrick Meir, Aaron M. Ellison

**Affiliations:** Programa de Pós-graduação em Zoologia, Instituto de Ciências Biológicas, Universidade Federal do Pará, Rua Augusto Corrêa, N° 1, 66075-110, Belém, Pará, Brazil; Coordenação de Ciências da Terra e Ecologia, Museu Paraense Emílio Goeldi, Av. Perimetral, N° 1901, 66077-830, Belém, Pará, Brazil; Instituto de Geociências, Universidade Federal do Pará, Rua Augusto Corrêa, N° 1, 66075-110, Belém, Pará, Brazil; Coordenação de Botânica, Museu Paraense Emílio Goeldi, Av. Perimetral, N° 1901, 66077-830, Belém, Pará, Brazil; Research School of Biology, Australian National University, Canberra ACT 2601, Australia; School of Geosciences, University of Edinburgh, Edinburgh EH93FF UK; Harvard Forest, Harvard University, Petersham, Massachusetts, USA

**Keywords:** Amazonia, climatic change, ESECAFLOR, Formicidae, tropical rainforest, water stress

## Abstract

Environmental change scenarios caused by low precipitation forecast species loss in tropical regions. We use one year of data from a long-term rainwater exclusion experiment in primary Amazonian rainforest to test whether induced water stress and covarying changes in soil moisture, soil respiration, tree species richness, diversity, size, and total biomass affected species richness and composition (relative abundance) of ground-dwelling ants. Induced drought reduced ant richness, whereas increased soil moisture and variability in tree biomass increased it. Species composition differed between control and rainfall-excluded plots. Occurrence of many ant species was strongly reduced by induced drought, but some generalist groups of ants were favored by it. The expected loss of ant species and changes in ant species composition in tropical forests likely will lead to cascading effects on ecosystem processes and the services they mediate.

## Introduction

Biological responses to ongoing climatic change are determined by organisms’ physiology, behavior, ecology, and evolutionary history (Diamond et al. 2016). Species inhabiting tropical forests have been considered physiologically more sensitive to climatic change because they are closer to their thermal tolerance limits than are organisms living in temperate environments (Diamond et al. 2016, Warren et al. 2018). Different scenarios of climatic change for the Amazon Basin include changes in temperature, precipitation, and relative humidity (Phillips et al. 2009, Davidson et al. 2012, Mora et al. 2013), and simulations forecast insect species losses by the year 2100 ranging from 6% with 1.5 °C warming to 49% with ~3.2 °C warming (Warren et al. 2018).

Many climate-change scenarios forecast droughts in the Amazon, but their effects on invertebrate biodiversity are virtually unknown (Ruivo et al. 2007, Diamond et al. 2012, Baccaro et al. 2013, Sánchez-Bayo and Wyckhuys 2019). Drought reduces photosynthesis and tends to kill larger trees (Meir et al. 2015, Costa et al. 2017). The associated loss of biomass and plant litter input will alter soil microclimate, which may reduce or eliminate foraging and nesting sites for ground-dwelling invertebrates (Byrne 1994, Fernandes et al. 2019). Loss of invertebrate species will also alter ecosystem processes and functions, including reduction of organic-matter decomposition and nutrient cycling rates and retention of soil humidity (Griffiths et al. 2018).

Ants interact with many plants and animals and are an important component of most terrestrial ecosystems (Hölldobler and Wilson 1990). Ants include thermophilic groups and are frequently used in global analyses on relationships between invertebrate diversity and climate (Gibb et al. 2015, Sánchez-Bayo and Wyckhuys 2019). Drought and increased temperature can increase abundance of thermophilic species while reducing resilience of ant assemblages (Folgarait 1998, Vasconcelos et al. 2010, Baccaro et al. 2013, Diamond et al. 2016). Ant species richness also tends to be lower in environments with low structural heterogeneity complexity, which also favor generalist species (Leal et al. 2012, Baccaro et al. 2013, Fernandes et al. 2019). Here, we test whether environmental and forest-structural changes caused by an experimentally induced drought in an eastern Amazonian rainforest alters species richness and composition of ground-dwelling ants. These data were collected as part of the long-term “Effects of forest drought” experiment (ESECAFLOR) (Meir et al. 2018).

## Materials and methods

### Study area

ESECAFLOR is a large-scale, long-term rainfall-exclusion experiment established in 2001 to study impacts of sustained low water supply (induced drought) on local microclimate, vegetation, and ecosystem processes in a *terra firme* rainforest (Meir et al. 2018). It is located at the Ferreira Penna Scientific Station (ECFPn) in the Caxiuanã National Forest (FLONA), Pará, Brazil (1.718° S, 51.46° W). Average air temperature at ECFPn ≈ 26 °C and daily variation in temperature < 4 °C. The *terra firme* forest on yellow oxisol soils is 10–15 m above river level. Average annual precipitation is 2,000–2,500 mm but < 100 mm/month falls during the dry season (June to November; Ruivo and Cunha 2003).

ESECAFLOR includes a pair of 1-ha (100×100-m) plots. In January 2002, plastic panels were installed 2 m above the soil in the **experimental plot** to exclude ≈ 50% of precipitable water (Appendix S1: Fig. S1A). This reduction is similar to the long-term forecast for the region (Marengo et al. 2018). Panels are maintained regularly; accumulated plant litter is removed and spread evenly on the ground below each panel. Fifty m away from the experimental plot, an equivalent 1-ha **control plot**, whose topography, soil type, and vegetation structure were similar to that of the experimental plot, was demarcated (Fisher et al. 2007, Meir et al. 2018).

### Ant sampling

The two ESECAFLOR plots were each subdivided into 25 20×20-m subplots. Ants were sampled using pitfall traps—an efficient method for collecting ground-dwelling ants in the Amazon (Souza et al. 2012)—in all subplots in October and November 2011 and again in February, March, May, July, August and September 2012 (Appendix S1: Fig. S1B). For pitfall traps, we used 500-ml cups (10-cm diameter) containing water, salt, and drops of detergent. The cups were covered to avoid overflow from rains and to avoid trapping larger arthropods and small vertebrates (Bestelmeyer et al. 2000). We placed four pitfall traps in each subplot in each month (total sampling effort = 1600 pitfall traps). Traps were left open in the field for 48 hours. Accumulated ants were then transferred into vials containing 80% ethanol; individuals from each species in each sample were mounted and identified. Voucher specimens were deposited in the Entomological Collection of the Museu Paraense Emílio Goeldi, Belém, Brazil.

### Data analysis

Rainfall exclusion altered the environmental variables in the ESECAFLOR experimental plot (Appendix S1: Table S1). We analyzed the effects of these drought-related changes in the forest environment and vegetation on ant species richness and composition in both space and time. For the “spatial model”, the 20×20-m subplots were considered as the sampling units (all data pooled over time); treating subplots as independent spatial replicates was justified because there was no spatial autocorrelation in the values of species richness among subplots within each 1-ha ESECAFLOR plot (control plot: Moran’s I = 0.035, *P* = 0.41; experimental plot: Moran’s I = 0.085; *P* = 0.93). For the “temporal model”, months were considered as the sampling units (all data in each of the two ESECAFLOR 1-ha plots pooled for each month). There was no significant collinearity among predictors in either the spatial or temporal models (Appendix S1: Figs. S2, S3).

To test experimental effects on species richness in both the spatial and temporal models, we used generalized linear mixed models (GLMMs) implemented in the “lme” function of the *nlme* package (Pinheiro et al. 2020) in the R software system (version 4.0.0; R Core Team 2020). Model selection was done using AICc values (Burnham and Anderson 2002) computed with the “dredge” function of the *MuMIn* package (Barton 2020). Model variables were included when their importance factors > 0.7 (on a 0–1 scale) based on their occurrence in the selected feasible models (Burnham and Anderson 2002).

To test whether ant species composition differed between treatments, we used Permutational Multivariate Analysis of Variance (PERMANOVA; Clarke, 1993) as implemented in the “adonis” function in the *vegan* package (Oksanen et al. 2019) with 9999 randomizations. Redundancy Analysis (RDA) was used to visualize ordinations using the “rda” function of the package *vegan* (Oksanen et al. 2019); the Bray-Curtis index was used as the distance measure in the ordinations (Gotelli et al. 2011). Analyses were included in the models by order of importance.

Finally, we classified species as specialists in the control and experimental plots using a multinomial Species Classification Method (CLAM) as implemented in the “clamtest” function of the *vegan* package (Oksanen et al. 2019). This test places species in four groups based on their occurrence frequency, allowing for a robust statistical classification with no need to exclude rare species (Chazdon et al. 2011). The four groups were: (i) preferentially occurring at the control plot; (ii) preferentially occurring at the experimental plot; (iii) no preference between categories or generalists; and (iv) species with a low sampling number or uncommon in the study area, unable to confidently attribute any of the three previous classifications. We used a predefined specialization value = 66%.

### Data and code availability

All data and R code are available from the Harvard Forest Data Archive and the Environmental Data Initiative (https://harvardforest1.fas.harvard.edu/exist/apps/datasets/showData.html?id=HF346).

## Results

In the two ESECAFLOR plots, we collected 217 ground-dwelling ant species, belonging to 54 genera and 8 subfamilies (Appendix S1: Table S2). We recorded 172 ant species in the control plot, and 157 species in the experimental plot; 112 species were recorded in both plots. The richest genus was *Pheidole* (43 species), followed by *Neivamyrmex*, *Gnamptogenys* and *Strumigenys* (10 species each).

The data supported our hypothesis that induced drought reduces ant species richness in both time and space. The model that best explained temporal changes in ant species richness included experimental treatment (plot), litter diversity, and their interaction (Fig. 1A, B; Appendix S1: Table S3). Fourteen spatial models had ΔAICc values < 2 (Appendix S1: Table S3), and variously included experimental treatment, litter diversity, treatment × litter diversity (Fig. 1C, D), and tree species richness (Appendix S1: Fig. S4).

**Fig. 1.**
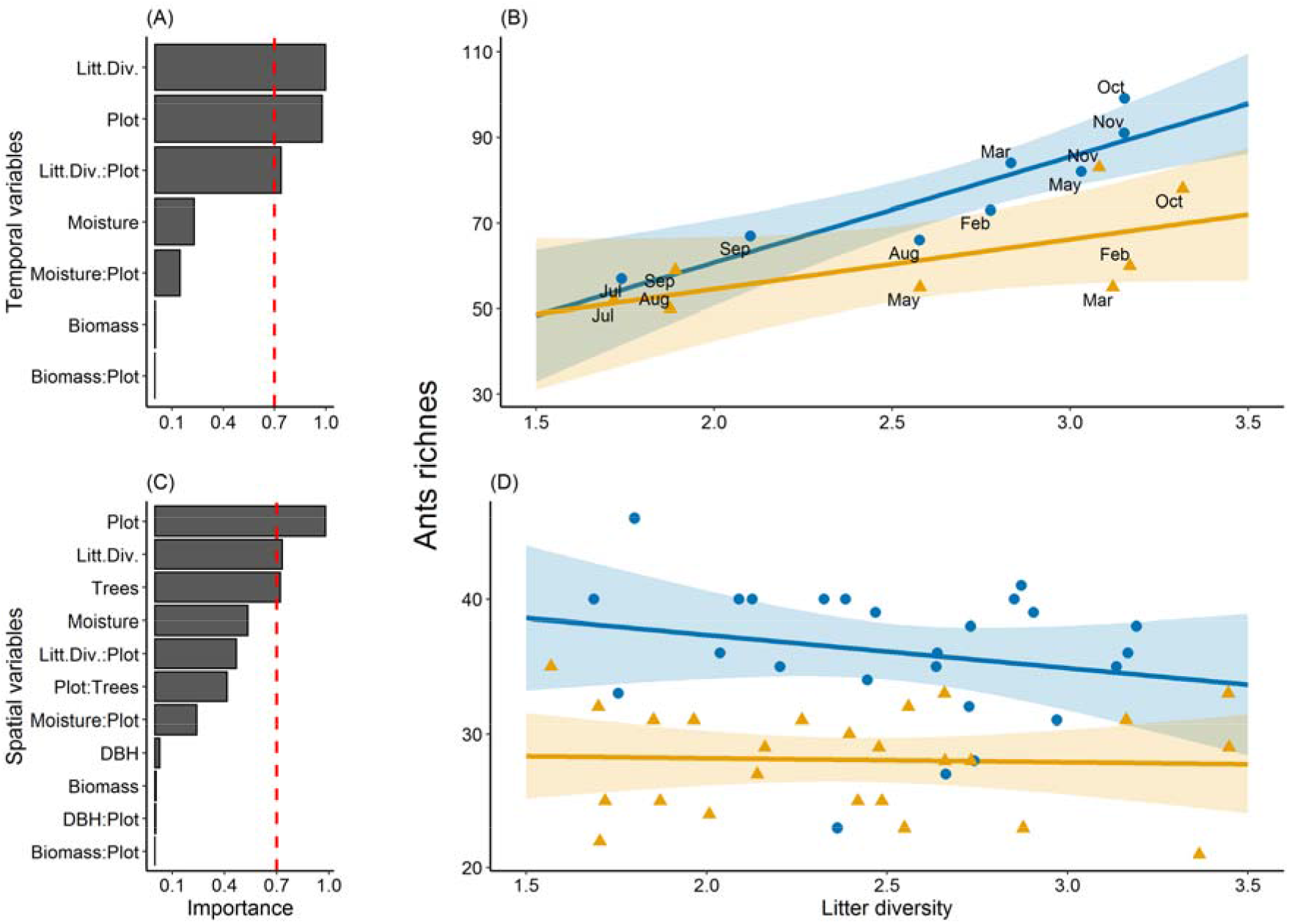
The relative importance of each variable and the relationship between variability in tree biomass and ant species richness in the (A, B) temporal and (C, D) spatial models. Variables whose relative importance > 0.7 (vertical red lines in A and C) were included in the models. In B and D, blue represents the control plot and yellow the experimental plot where rainfall was excluded. (Abbreviations; Litt.Div. = Litter diversity, Resp. = Respiration, DBH = Diameter at breast height).

The data also supported our hypothesis that ant species composition differed in control and drought-induced conditions (Appendix S1: Table S4). Species composition was affected by experimental treatment, soil moisture (spatial and temporal models), and litter diversity (temporal model only) (Fig. 2A; Appendix S1: Table S4). For the temporal model, RDA explained 29% of the total variation on the first two axes (Appendix S1: Table S5). Moisture had the strongest contribution to axis 1 and was associated with the species group in the control plot (Fig. 2A). Litter diversity had the strongest contribution to axis 2 (Fig. 2A). For the spatial model, the first two axes of the RDA explained < 10% of the total variation in the data (Fig. 2B). Soil moisture again loaded most strongly on axis 1 (Fig. 2B; Appendix S1: Table S5).

**Fig. 2.**
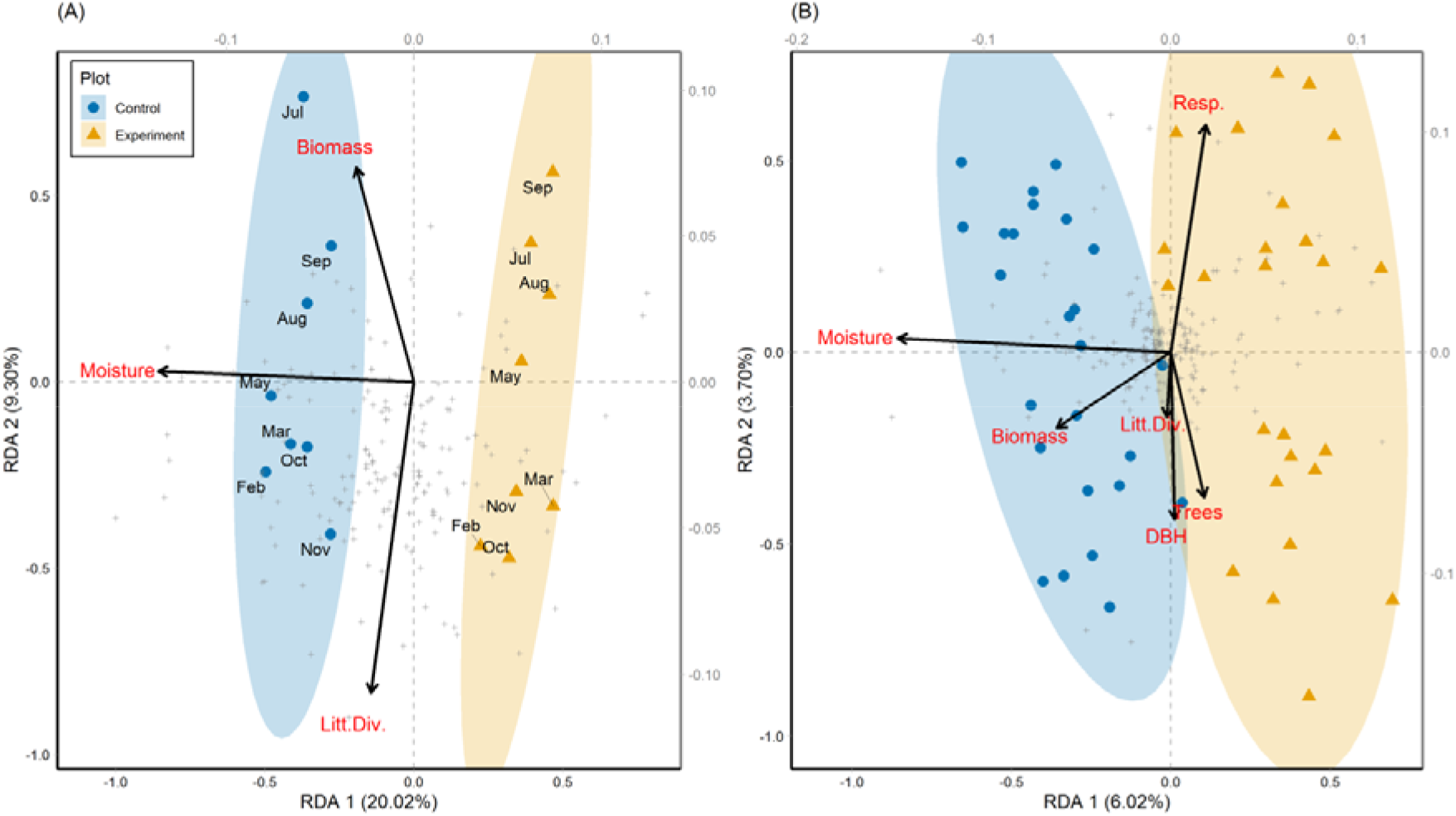
RDA results for the (A) temporal and (B) spatial models of ant assemblages in the control (blue) and experimental (yellow) plots (statistics in Appendix S1: Tables S4, S5). Abbreviated variable names as in Fig. 1. Grey points represent the locations in ordination space of the 217 ant species collected.

Finally, we found specialist species in both the experimental and control plots (Fig. 3). Multinomial classification identified 12 specialists in the control plot (*Neoponera verenae*, *Ochetomyrmex neopolitus*, *Oc. semipolitus*, *Odontomachus* nr. *bauri*, *Od.* nr. *hematodus*, *Octostruma betschi*, *Pheidole cursor*, *P. jeannei*, *Pheidole* in the *Fallax* group and another unidentified, *Strumigenys denticulata* and *Solenopsis* nr. *virulens*) and 12 species in induced-drought conditions (*Apterostigma* nr. *pilosum*, *Mayaponera constricta*, *M. arhuaca*, *Neivamyrmex pseudops*, *Pachycondyla crassinoda*, *Paratrachymyrmex* nr. *bugnioni*, *Pheidole bruesi*, *P. triconstricta*, and an unnamed species of each of *Azteca*, *Crematogaster*, *Neivamyrmex*, and *Nylanderia)*. Of the remaining species, 44 (20%) showed no preference among plots and 149 (69%) occurred at such low frequencies that plot preferences could not be determined (Fig. 3).

**Fig. 3.**
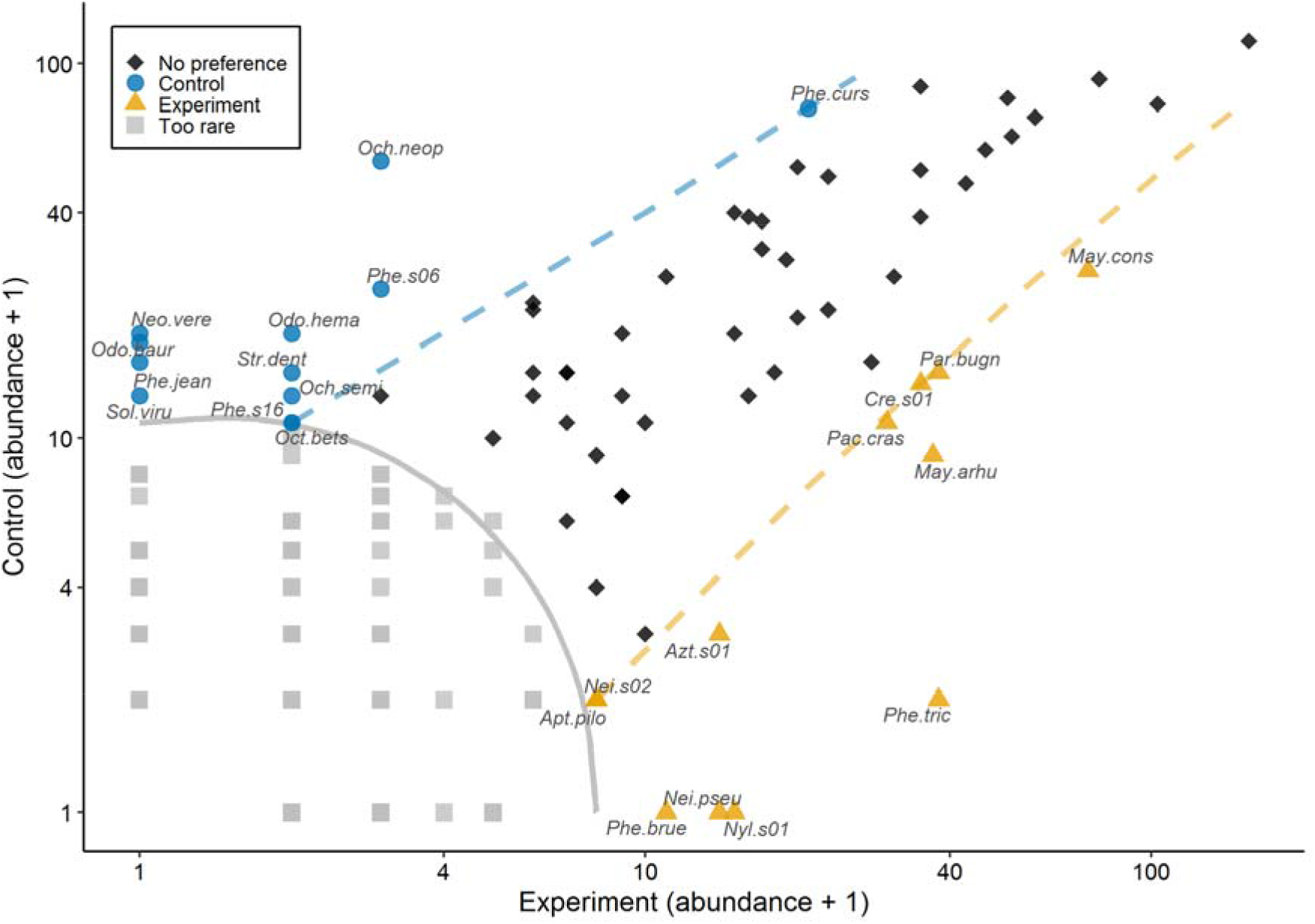
CLAM analysis classifying the 217 collected ant species into four classes based on their frequency of occurrence in the control and experimental plots: no preference or generalists (black diamonds); too rare to be assigned a specificity preference (gray squares); associated with the control plot (i.e., natural forest) (blue circles): associated with the experimental plot (i.e., drought-induced forest) (yellow triangles). Species codes given in Appendix S1: Table S2.

## Discussion

We have documented for the first time the effects of drought induced by rainwater exclusion on ant assembles in a primary Amazonian rainforest. Based on one year of sampling period in a long-term, large-scale rainfall-exclusion (induced-drought) experiment, our results strongly suggest that ant richness and composition are influenced by drought in the forest and are coupled with changes in both soil moisture content and litter diversity. We also identified some specialist ant species in a “natural” rainforest plot and in an adjacent, drier forest plot where rainfall was experimentally excluded.

### Species richness

Species richness was lower in the plot where rainfall was excluded. At 25-km^2^ and basin-wide scales, humid Amazonian forests have higher ant species richness (Vasconcelos et al. 2010, Baccaro et al. 2013). In tropical forests, ants have different strategies to avoid desiccation, such as foraging at night or within plant litter (Kaspari and Weiser 2000). However, the combination of high temperatures and reduced precipitation may reduce the survival or competitive ability of some tropical forest species (Debinski et al. 2013, Diamond et al. 2016).

### Species composition

Species composition differed between the 1-ha control plot and the 1-ha plot from which ≈50% of the rainfall was excluded. Compositional changes as a function of drought induced by climatic change have been reported previously for butterflies (Carnicer et al. 2019) and amphibians (Gouveia and Correia 2016). Ant assemblages are known to change with increasing temperature (Diamond et al. 2012, 2016, Menke et al. 2014), which in turn can reduce soil moisture content. The loss of soil moisture resulting from reduction in precipitation, seasonal flooding, or deeper groundwater also can directly affect ant assemblages (Vasconcelos et al. 2010, Baccaro et al. 2013). The reduction of water availability in tropical forests also can affect ants indirectly through their effects on trees. For example, a lower diversity of items in the litter can affect the taxonomic and functional diversity of ant communities (Folgarait 1998, Coyle et al. 2017), with long-term effects on environmental recovery (Martello et al. 2018).

### Species associations

The identity of ant species that are associated with reduction in rainfall can help inform our understanding of how drought may alter ant community assembly structure. For instance, fungus-growing ants (*Apterostigma* and *Paratrachymyrmex* spp.) were associated with the drier (experimental) conditions. These genera also tend to increase in frequency in fragmented or simplified environments (Ribas et al. 2012, Baccaro et al. 2013). Their drought tolerance is related to their nesting under trunks or in the ground rather than directly on the leaf-litter. Indeed, fauna living in deeper soil may be more resilient to drought or drying than more superficial dwellers (Coyle et al. 2017). Further, among the 23 large-sized epigeic predator species in our study (Silva and Brandão 2014), only three (*Mayaponera constricta, M. arhuaca* and *Pachycondyla crassinoda*) were more common in the experimental (drought-induced) plot. These three species were only found in the experimental plot, and *P. crassinoda* is associated with drier habitats in the Amazon (Baccaro et al. 2013). Overall, our data clearly identify generalist, thermophilic species associated with simplified environments when drought was induced in an otherwise wet tropical forest. Other large ant species (especially *Odontomachus* and *Neoponera* spp) were associated with the control plot. Exclusion of large ants because of increased temperatures has been recorded in a recent global study (Gibb et al. 2018). Some of these may be important seed dispersers, but few studies have evaluated ant-mediated seed dispersal in the Amazon.

## Conclusion

Water stress affects ant assemblages, reducing richness and altering species composition, mainly through the increased occurrence of thermophilic species. When throughfall was experimentally reduced in an Amazonian forest, consequent effects of variability in biomass and soil moisture were the most important predictors of species richness and composition of ground-dwelling ant species. Changes caused by droughts may have direct effects on soil fauna that depend on organic matter on the soil surface, and indirect effects that cascade through trees that structure the forest. Our results were obtained from 2 ha of experimental rainforest but extended El Niño events or climatic changes may cause similar effects at much larger scales throughout Amazonia.

## Acknowledgments

The authors were supported by grants from Conselho Nacional de Desenvolvimento Científico e Tecnológico (CNPq: RPSA; PCI-DC-313516/2015-4 and AME; PCI-170220/2016-8, PCI-170088/2017-0). This study was financed in part by the Coordenação de Aperfeiçoamento de Pessoal de Nível Superior - Brasil (CAPES) - Finance Code 001. We warmly thank all of the staff of LBA/ ESECAFLOR and Estação Científica Ferreira Penna for logistical support and accommodation. We are grateful to Arleu Viana-Jr, Fabrício Baccaro, Helder Espírito-Santo, Joudelys Andrade-Silva, Leandro Juen and Raphael Ligeiro for ideas and suggestions in analysis. Adrian Troya, Alexandre Ferreira, Emília Albuquerque, Joudellys Andrade-Silva, Júlio Chaul, Lívia Prado, Otávio Silva, Rodrigo Feitosa and Thiago Silva for assisting with ant identification.

