## Appendix S1 for "Induced drought strongly affects richness and composition of ground-dwelling ants in the eastern Amazon"

*Environmental and vegetation data at ESECAFLOR*

Researchers involved in the ESECAFLOR experiment (Fig. S1) collect data that describe the dynamics of both environment and vegetation (e.g., solar radiation/min, air temperature and humidity at 1 and 10 m above ground/min, ground temperature and soil moisture at 5-, 10- and 50-cm depth/min, precipitation/hour). For the analyses presented here, we used variables related to the food resources and the diversity of microhabitats to ant foraging and nesting to test the relationship between ant assemblages and the environment (Byrne 1994, Kaspari and Weiser 2000, Vasconcelos et al. 2010, Jenkins et al. 2011, Baccaro et al. 2013, Diamond et al. 2016, Coyle et al. 2017). Furthermore, we selected those variables that match both the period of ant sampling (temporal data) and the spatial data (20 × 20 subplots). Thus, we selected three covariates for temporal analysis, and six covariates for spatial analyses (Table S1). Variables used were not collinear (Figs. S2, S3).

Soil moisture is recorded every hour by a weather station established at the center of each plot (subplot 13; Figure S1b) using sensors (*CS616; Campbell Scientific*) installed at ground level, placed on the right and left sides of the tower. We used the average of all data collected each month. More information can be found in Fisher et al. (2007).

Plant litter measurements are made in each parcel using 25 1-m$^2$ plant-litter traps in each subplot collected once each month (Fig. S1). Collectors are installed at 1 m above ground at the control plot and above plastic panels at the experimental plot. Litter is collected from all traps and sorted into leaf material, reproductive material (flowers and fruits), woody material <2 cm in diameter (twigs) and non-identifiable plant material, which comprised only a small fraction of annual litterfall, on average (<5%). Following separation, the material is then dried to a constant temperature. Here we use total biomass, or the sum of all collected material, as a variable. We calculated the litter diversity using Hill number in order 1 (Shannon diversity equivalent; Chao et al. 2014), based in biomass, in grams, of each separate component. We used the summed biomass in the plots for each month. Details on biomass collection and time series are given in Rowland et al. (2018). Biomass and litter diversity were also used as spatial variables. For that purpose, we used the sum of all monthly biomass in each parcel.

Soil respiration is measured using static chambers. Chambers (polyvinyl chloride [PVC] rings measuring 10 × 10 cm [height × internal diameter]) are set ≈2 cm into the ground at the center of each subplot. $CO_2$ is measured by attaching a PPSystems Soil Respiration Chamber (model SRC 1) to the PVC ring; the chamber in turn is connected to a portable infrared gas analyzer (*Environment Gas Monitor* - EGM-5, PPSystems, USA). Respiration was measured in six out of the eight months (October and December 2010 and June - September 2011). We used the average monthly respiration (μmol $CO_2$ $m^{-2}$ $s^{-1}$) in each parcel.

Soil moisture also was measured gravimetrically during soil respiration measurements. The material was sampled using a soil auger (at 0-10 cm depth), near each respiration ring in each parcel. Soil samples were identified and stored in sealed plastic containers. Wet matter was weighed in the laboratory using a precision scale. Afterwards, samples were dried in an oven at 105 °C for 24 hours for dry matter weighing. Soil moisture content was calculated as the difference between wet and dry matter weights (Fisher et al. 2007). We used the average monthly soil moisture for each parcel.

In March 2010, all tree species with a diameter at breast height (DBH) above 10 cm were identified and had their DBH measured. We used tree richness and the average DBH (cm) for each 20 x 20 m subplot.

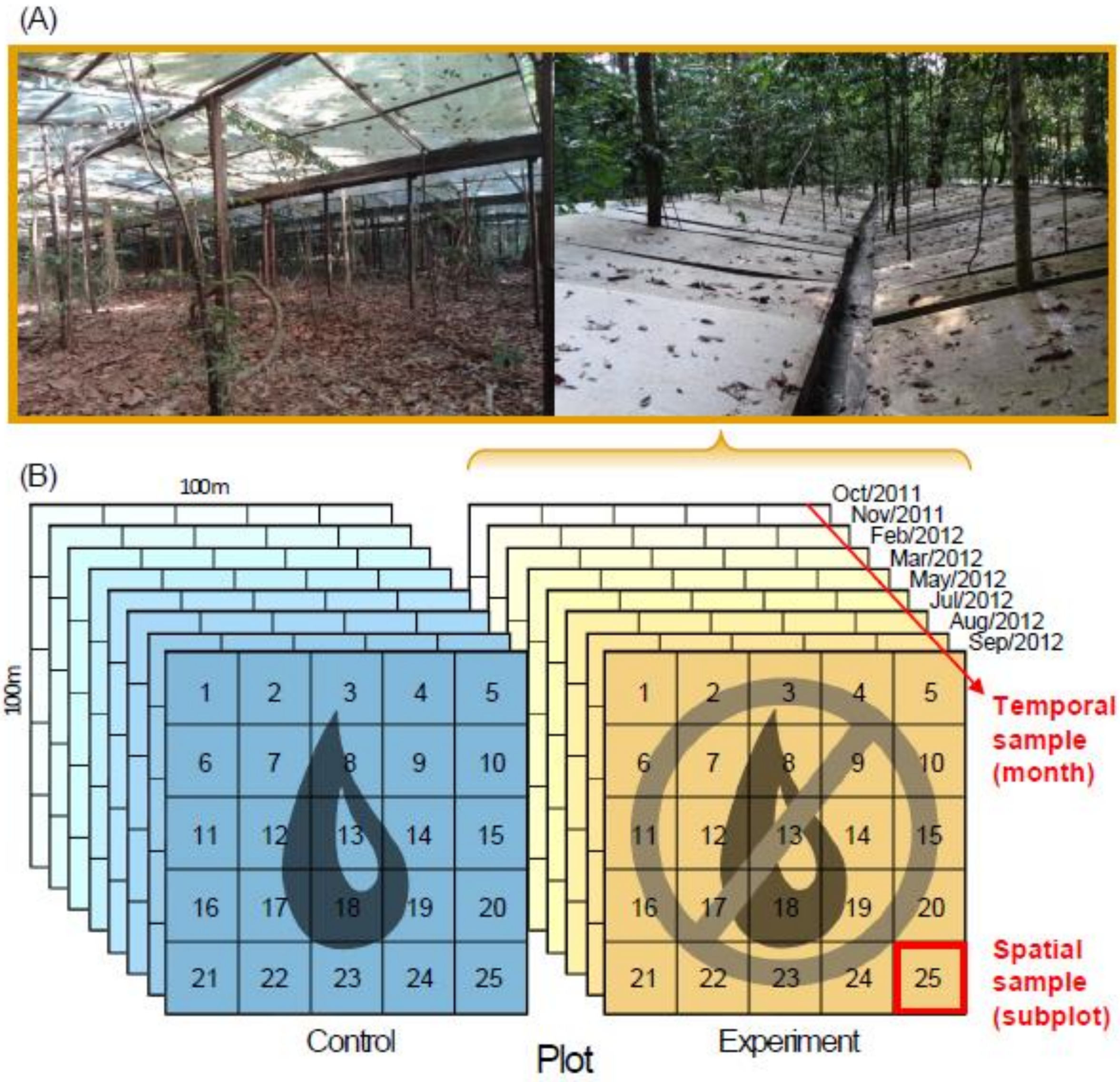


Fig. S1. (A) Photos obtained from below (left) and above (right) of the plastic panels that exclude rainwater at the experimental plot. (B) Experimental design for both plots (control and experimental), illustrating the temporal (months) and space (subplots) samples for collecting ants and environmental data, in ESECAFLOR, Ferreira Penna Scientific Station (ECFPn), Caxiuanã National Forest (FLONA), Pará, Brazil.

Table S1. Covariates selected for analyses of ant data in ESECAFLOR experiment, sample scale (temporal and spatial), and data dimension. The response variable in each model was species richness or species composition. The temporal data was defined as the eight months of sampling in each plot (N = 16). The spatial data was defined as the 25 subplots of 20 x 20 m in each plot (N = 50).

| **Predictor** | **Samples** | **Description** | **Data length** |
|---|---|---|---|
| **Moisture** | Spatial and Temporal | Soil moisture collected by the meteorological tower and manually in each plot. Measured in %. | Spatial: 50 x 400 Temporal: 8 x 23,424 |
| **Litter diversity** | Spatial and Temporal | Components of biomass (measured in grams: leaves, twigs, flowers, fruits, and miscellaneous) submitted Hill number in order 1. | Spatial and Temporal: 50 x 2,400 |
| **Biomass** | Spatial and Temporal | Sum of all biomass items. Measured in grams. | Spatial and Temporal: 50 x 400 |
| **Respiration** | Spatial | Amount of $CO_2$ released. Measured in μmol $CO_2$ $m^{-2}$ $s^{-1}$. | 50 x 400 |
| **Trees** | Spatial | Richness of tree species over 10 cm in diameter at breast height per plot. Measured in number of species. | 50 x 266 tree species |
| **DBH** | Spatial | Mean diameter at breast height (DBH), per plot, of individuals over 10 cm in diameter. Measured in centimeters. | 1040 individuals |

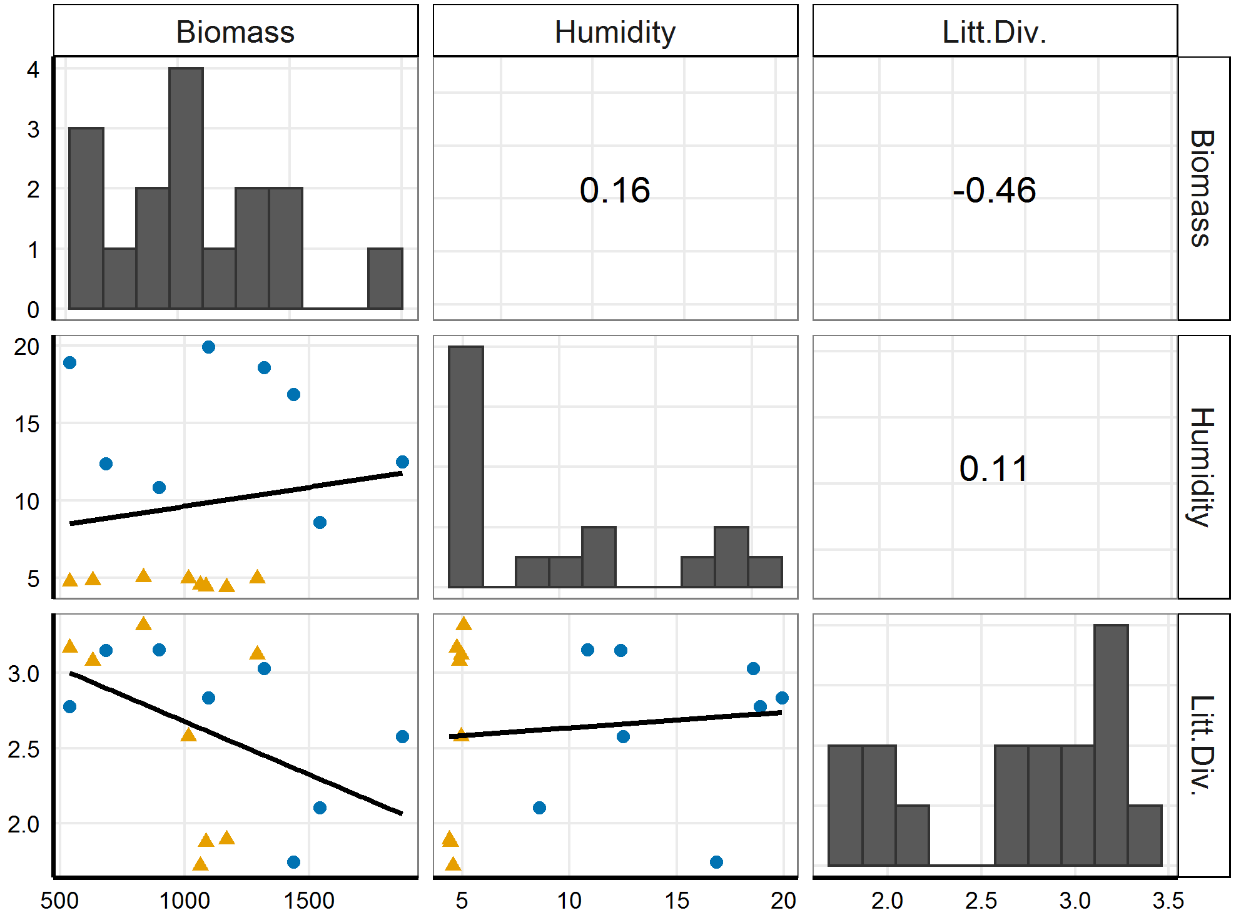


Fig. S2. Correlation panel between predictor variables in the temporal model for the control and experimental plots in the ESECAFLOR experiment. Blue circles = control plot; yellow triangles = experimental plot; the black line shows the trend in the relationship. (Litt.Div. = Litter diversity).

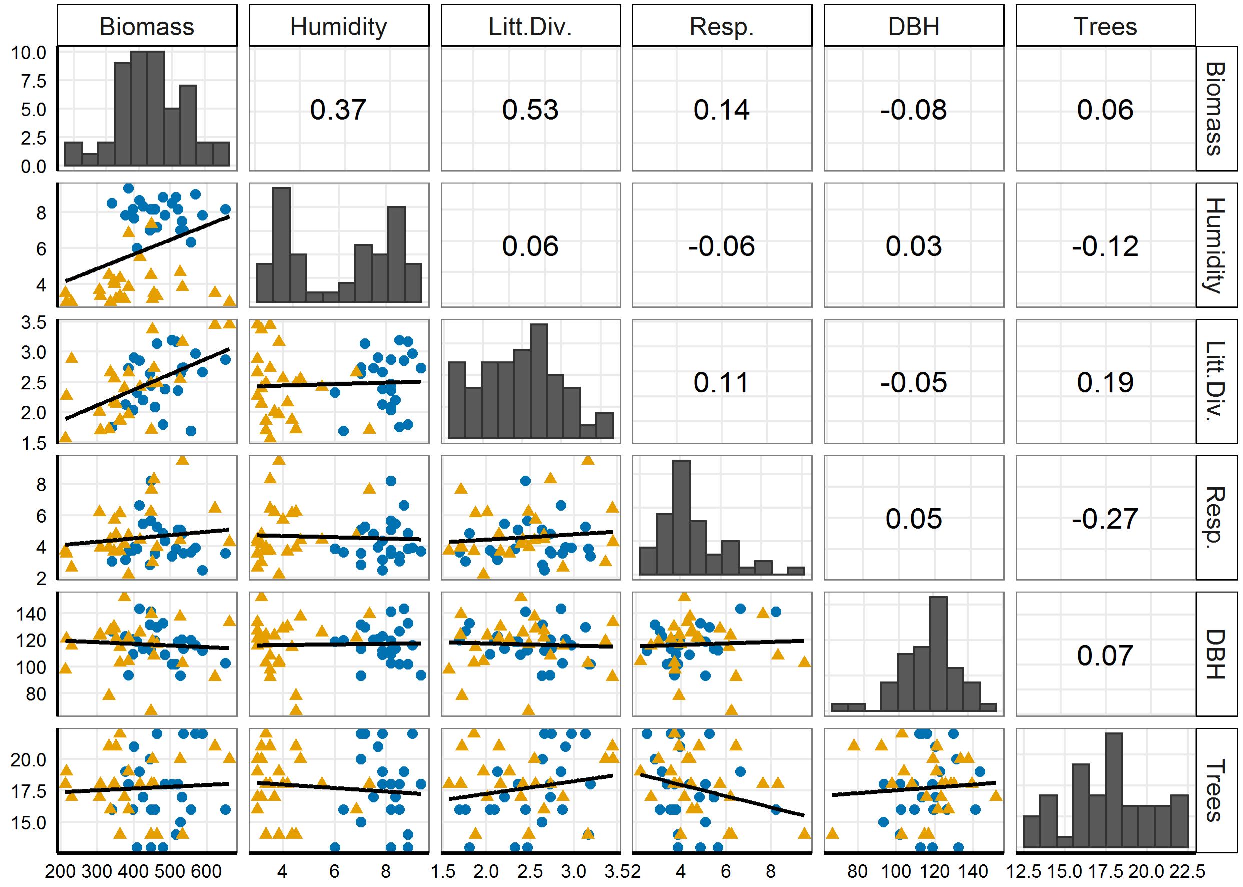


Fig. S3. Correlation panel between predictor variables in the spatial model for the control and experimental plots in the ESECAFLOR experiment. Blue circles = control plot; yellow triangles = experimental plot; the black line shows the trend in the relationship. (Litt.Div. = Litter diversity, Resp. = Respiration).

Table S2. Frequency of occurrence of ant species collected at the control plot (**Con**) and experimental plot (**Exp**) in the ESECAFLOR project, Floresta Nacional de Caxiuanã, Pará, Brazil, in 2011 and 2012. (Records: subplots 0-25; months 0-8; total 0-200).

| | | **Occurrences** | | | | | |
|---|---|---|---|---|---|---|---|
| | | **Subplots** | | **Months** | | **Total** | |
| **Subfamily or species** | **Abbr.** | **Con** | **Exp** | **Con** | **Exp** | **Con** | **Exp** |
| **Dolichoderinae** | | | | | | | |
| *Azteca* MPEG 01 | Azt.s01 | 2 | 6 | 2 | 8 | 2 | 13 |
| *Azteca* MPEG 02 | Azt.s02 | 6 | 6 | 3 | 5 | 6 | 8 |
| *Azteca* MPEG 03 | Azt.s03 | 2 | 7 | 2 | 5 | 2 | 9 |
| *Azteca* MPEG 04 | Azt.s04 | 0 | 1 | 0 | 1 | 0 | 1 |
| *Azteca* MPEG 05 | Azt.s05 | 2 | 0 | 2 | 0 | 2 | 0 |
| *Dolichoderus attelaboides* (Fabricius, 1775) | Dol.atte | 1 | 2 | 1 | 1 | 1 | 2 |
| *Dolichoderus bispinosus* (Olivier, 1792) | Dol.bisp | 13 | 11 | 8 | 6 | 29 | 18 |
| *Dolichoderus decollatus* Smith, 1858 | Dol.deco | 5 | 3 | 6 | 3 | 6 | 3 |
| *Dolichoderus gagates* Emery, 1890 | Dol.gaga | 1 | 0 | 1 | 0 | 1 | 0 |
| *Dolichoderus imitator* Emery, 1894 | Dol.imit | 1 | 0 | 1 | 0 | 1 | 0 |
| *Dolichoderus laminatus* (Mayr, 1870) | Dol.lami | 0 | 1 | 0 | 1 | 0 | 1 |
| *Dolichoderus lutosus* (Smith, 1858) | Dol.luto | 2 | 1 | 2 | 1 | 2 | 1 |
| *Dolichoderus quadridenticulatus* (Roger, 1862) | Dol.quad | 2 | 0 | 2 | 0 | 2 | 0 |
| *Dolichoderus* MPEG 03 | Dol.s01 | 1 | 0 | 1 | 0 | 1 | 0 |
| **Dorylinae** | | | | | | | |
| *Cylindromyrmex striatus* Mayr, 1870 | Cyl.stri | 1 | 0 | 1 | 0 | 1 | 0 |
| *Eciton* nr. *burchellii* | Eci.burc | 5 | 1 | 3 | 1 | 5 | 1 |
| *Eciton* nr. *drepanophorum* | Eci.drep | 2 | 1 | 1 | 1 | 2 | 2 |
| *Eciton* nr. *mexicanum* | Eci.mexi | 6 | 2 | 3 | 1 | 6 | 2 |
| *Eciton rapax* Smith, 1855 | Eci.rapa | 3 | 0 | 1 | 0 | 4 | 0 |

| | | | | | | | |
|---|---|---|---|---|---|---|---|
| *Labidus coecus* (Latreille, 1802) | Lab.coec | 14 | 17 | 5 | 6 | 15 | 27 |
| *Labidus praedator* (Smith, 1858) | Lab.prae | 7 | 4 | 1 | 2 | 9 | 4 |
| *Labidus spininodis* (Emery, 1890) | Lab.spin | 20 | 10 | 6 | 3 | 38 | 15 |
| *Neivamyrmex cristatus* (André, 1889) | Nei.cris | 0 | 2 | 0 | 1 | 0 | 2 |
| *Neivamyrmex pilosus* (Smith, 1858) | Nei.pilo | 1 | 5 | 1 | 4 | 1 | 5 |
| *Neivamyrmex pseudops* (Forel, 1909) | Nei.pseu | 0 | 10 | 0 | 2 | 0 | 13 |
| *Neivamyrmex* nr. *swainsonii* | Nei.swai | 1 | 0 | 1 | 0 | 1 | 0 |
| *Neivamyrmex* MPEG 01 | Nei.s01 | 1 | 0 | 1 | 0 | 1 | 0 |
| *Neivamyrmex* MPEG 02 | Nei.s02 | 1 | 7 | 1 | 3 | 1 | 7 |
| *Neivamyrmex* MPEG 03 | Nei.s03 | 1 | 0 | 1 | 0 | 1 | 0 |
| *Neivamyrmex* MPEG 04 | Nei.s04 | 2 | 1 | 2 | 1 | 2 | 1 |
| *Neivamyrmex* MPEG 05 | Nei.s05 | 0 | 2 | 0 | 2 | 0 | 2 |
| *Neivamyrmex* MPEG 08 | Nei.s06 | 1 | 0 | 1 | 0 | 1 | 0 |
| *Nomamyrmex esenbeckii* (Westwood, 1842) | Nom.esen | 5 | 2 | 3 | 2 | 7 | 2 |
| *Nomamyrmex hartigii* (Westwood, 1842) | Nom.hart | 1 | 0 | 1 | 0 | 1 | 0 |
| **Ectatomminae** | | | | | | | |
| *Ectatomma edentatum* Roger, 1863 | Ect.eden | 10 | 4 | 7 | 7 | 18 | 14 |
| *Ectatomma lugens* Emery, 1894 | Ect.luge | 3 | 2 | 5 | 4 | 5 | 4 |
| *Ectatomma tuberculatum* (Olivier, 1792) | Ect.tube | 3 | 0 | 4 | 0 | 4 | 0 |
| *Gnamptogenys concinna* (Smith, 1858) | Gna.conc | 1 | 0 | 1 | 0 | 1 | 0 |
| *Gnamptogenys horni* (Santschi, 1929) | Gna.horn | 24 | 16 | 8 | 7 | 86 | 34 |
| *Gnamptogenys kempfi* Lenko, 1964 | Gna.kemp | 2 | 1 | 2 | 1 | 2 | 1 |
| *Gnamptogenys pleurodon* (Emery, 1896) | Gna.pleu | 1 | 0 | 1 | 0 | 1 | 0 |
| *Gnamptogenys sulcata* (Smith, 1858) | Gna.sulc | 4 | 1 | 3 | 1 | 4 | 1 |
| *Gnamptogenys tortuolosa* (Smith, 1858) | Gna.tort | 10 | 3 | 6 | 2 | 14 | 5 |

| | | | | | | | |
|---|---|---|---|---|---|---|---|
| *Gnamptogenys* nr. *curvoclypeata* | Gna.curv | 1 | 0 | 1 | 0 | 1 | 0 |
| *Gnamptogenys* nr. *rastrata* | Gna.rast | 2 | 0 | 2 | 0 | 2 | 0 |
| *Gnamptogenys* MPEG 02 | Gna.s01 | 0 | 4 | 0 | 4 | 0 | 4 |
| *Gnamptogenys* MPEG 03 | Gna.s02 | 0 | 1 | 0 | 1 | 0 | 1 |
| **Formicinae** | | | | | | | |
| *Brachymyrmex* MPEG 01 | Bra.s01 | 3 | 1 | 3 | 2 | 5 | 2 |
| *Brachymyrmex* MPEG 02 | Bra.s02 | 3 | 1 | 4 | 1 | 5 | 1 |
| *Brachymyrmex* MPEG 03 | Bra.s03 | 0 | 2 | 0 | 3 | 0 | 4 |
| *Brachymyrmex* MPEG 04 | Bra.s04 | 0 | 1 | 0 | 1 | 0 | 1 |
| *Camponotus femoratus* (Fabricius, 1804) | Cam.femo | 11 | 8 | 8 | 8 | 39 | 14 |
| *Camponotus mucronatus* Emery, 1890 | Cam.mucr | 1 | 0 | 1 | 0 | 1 | 0 |
| *Camponotus* nr. *melatoticus* | Cam.mela | 0 | 2 | 0 | 2 | 0 | 2 |
| *Camponotus* MPEG 01 | Cam.s01 | 1 | 1 | 1 | 1 | 1 | 1 |
| *Camponotus* MPEG 02 | Cam.s02 | 0 | 1 | 0 | 1 | 0 | 1 |
| *Gigantiops destructor* (Fabricius, 1804) | Gig.dest | 1 | 0 | 1 | 0 | 1 | 0 |
| *Myrmelachista* nr. *bambusarum* | Myr.bamb | 0 | 1 | 0 | 1 | 0 | 1 |
| *Nylanderia* MPEG 01 | Nyl.s01 | 0 | 6 | 0 | 5 | 0 | 14 |
| *Nylanderia* MPEG 02 | Nyl.s02 | 10 | 12 | 6 | 7 | 12 | 15 |
| *Nylanderia* MPEG 03 | Nyl.s03 | 5 | 1 | 4 | 1 | 8 | 1 |
| *Nylanderia* MPEG 04 | Nyl.s04 | 1 | 0 | 1 | 0 | 1 | 0 |
| *Nylanderia* MPEG 05 | Nyl.s05 | 0 | 2 | 0 | 3 | 0 | 3 |
| *Nylanderia* MPEG 06 | Nyl.s06 | 0 | 1 | 0 | 1 | 0 | 1 |
| *Nylanderia* MPEG 07 | Nyl.s07 | 0 | 1 | 0 | 1 | 0 | 1 |
| *Nylanderia* MPEG 08 | Nyl.s08 | 0 | 3 | 0 | 3 | 0 | 4 |
| Myrmicinae | | | | | | | |
| *Acromyrmex* MPEG 01 | Acr.s01 | 0 | 1 | 0 | 1 | 0 | 1 |
| *Acromyrmex* MPEG 02 | Acr.s02 | 1 | 2 | 1 | 1 | 1 | 2 |
| *Allomerus octoarticulatus* Mayr, 1878 | All.octo | 1 | 0 | 1 | 0 | 1 | 0 |

| | | | | | | | |
|---|---|---|---|---|---|---|---|
| *Apterostigma urichii* Forel, 1893 | Apt.uric | 0 | 1 | 0 | 1 | 0 | 1 |
| *Apterostigma* nr. *pilosum* | Apt.pilo | 1 | 4 | 1 | 4 | 1 | 7 |
| *Apterostigma* MPEG 02 | Apt.s01 | 0 | 1 | 0 | 1 | 0 | 1 |
| *Apterostigma* MPEG 03 | Apt.s02 | 0 | 1 | 0 | 1 | 0 | 1 |
| *Apterostigma* MPEG 04 | Apt.s03 | 1 | 0 | 1 | 0 | 1 | 0 |
| *Atta* MPEG 01 | Att.s04 | 0 | 1 | 0 | 2 | 0 | 2 |
| *Blepharidatta brasiliensis* Wheeler, 1915 | Ble.bras | 15 | 9 | 8 | 7 | 37 | 16 |
| *Carebara* (gr. lignata) MPEG 01 | Car.s01 | 2 | 0 | 2 | 0 | 3 | 0 |
| *Carebara* (gr. lignata) MPEG 02 | Car.s02 | 3 | 1 | 2 | 1 | 3 | 1 |
| *Carebara* MPEG 03 | Car.s03 | 0 | 1 | 0 | 1 | 0 | 1 |
| *Cephalotes atratus* (Linnaeus, 1758) | Cep.atra | 1 | 3 | 1 | 3 | 1 | 5 |
| *Cephalotes maculatus* (Smith, 1876) | Cep.macu | 0 | 1 | 0 | 1 | 0 | 1 |
| *Cephalotes* MPEG 03 | Cep.s01 | 0 | 1 | 0 | 1 | 0 | 1 |
| *Crematogaster brasiliensis* Mayr, 1878 | Cre.bras | 3 | 1 | 3 | 1 | 4 | 1 |
| *Crematogaster crinosa* Mayr, 1862 | Cre.crin | 3 | 4 | 3 | 4 | 4 | 4 |
| *Crematogaster flavosensitiva* Longino, 2003 | Cre.flav | 4 | 3 | 5 | 3 | 5 | 3 |
| *Crematogaster sotobosque* Longino, 2003 | Cre.soto | 17 | 13 | 8 | 8 | 47 | 42 |
| *Crematogaster* nr. *longispina* | Cre.long | 4 | 0 | 3 | 0 | 4 | 0 |
| *Crematogaster* MPEG 01 | Cre.s01 | 8 | 15 | 7 | 7 | 13 | 34 |
| *Crematogaster* MPEG 02 | Cre.s02 | 18 | 18 | 8 | 8 | 77 | 102 |
| *Crematogaster* MPEG 03 | Cre.s03 | 0 | 2 | 0 | 2 | 0 | 2 |
| *Crematogaster* MPEG 04 | Cre.s04 | 0 | 1 | 0 | 2 | 0 | 2 |
| *Cyphomyrmex laevigatus* Weber, 1938 | Cyp.laev | 4 | 1 | 5 | 1 | 5 | 1 |
| *Cyphomyrmex peltatus* Kempf, 1966 | Cyp.pelt | 11 | 5 | 6 | 4 | 14 | 6 |
| *Cyphomyrmex* nr. *minutus* | Cyp.minu | 3 | 4 | 3 | 4 | 3 | 4 |
| *Daceton armigerum* (Latreille, 1802) | Dac.armi | 1 | 2 | 1 | 2 | 1 | 2 |

| | | | | | | | |
|---|---|---|---|---|---|---|---|
| *Hylomyrma immanis* Kempf, 1973 | Hyl.imma | 0 | 2 | 0 | 2 | 0 | 2 |
| *Hylomyrma* MPEG 02 | Hyl.s01 | 0 | 1 | 0 | 1 | 0 | 1 |
| *Megalomyrmex cuatiara* Brandão, 1990 | Meg.cuat | 2 | 0 | 2 | 0 | 2 | 0 |
| *Megalomyrmex incisus* Smith, 1947 | Meg.inci | 1 | 1 | 1 | 1 | 1 | 1 |
| *Megalomyrmex* MPEG 01 | Meg.s01 | 1 | 0 | 1 | 0 | 1 | 0 |
| *Megalomyrmex* MPEG 02 | Meg.s02 | 1 | 0 | 1 | 0 | 1 | 0 |
| *Monomorium floricola* (Jerdon, 1851) | Mon.flor | 2 | 0 | 2 | 0 | 2 | 0 |
| *Mycetarotes* nr. *acutus* | Mya.acut | 1 | 0 | 2 | 0 | 2 | 0 |
| *Mycetomoellerius* nr. *farinosus* | Myl.fari | 1 | 3 | 1 | 2 | 1 | 3 |
| *Mycocepurus* nr. *smithii* | Myp.smit | 1 | 1 | 1 | 1 | 1 | 1 |
| *Myrmicocrypta* nr. *foreli* | Myr.fore | 4 | 2 | 3 | 2 | 5 | 2 |
| *Myrmicocrypta* MPEG 01 | Myr.s01 | 4 | 1 | 4 | 2 | 4 | 2 |
| *Myrmicocrypta* MPEG 03 | Myr.s02 | 1 | 1 | 1 | 1 | 1 | 1 |
| *Nesomyrmex* MPEG 01 | Nes.s03 | 0 | 1 | 0 | 1 | 0 | 1 |
| *Nesomyrmex* MPEG 02 | Nes.s04 | 0 | 1 | 0 | 1 | 0 | 1 |
| *Ochetomyrmex neopolitus* Fernández, 2003 | Och.neop | 13 | 2 | 8 | 2 | 54 | 2 |
| *Ochetomyrmex semipolitus* Mayr, 1878 | Och.semi | 8 | 1 | 5 | 1 | 12 | 1 |
| *Octostruma betschi* Perrault, 1988 | Oct.bets | 8 | 1 | 4 | 1 | 10 | 1 |
| *Octostruma iheringi* (Emery, 1888) | Oct.iher | 3 | 1 | 3 | 1 | 5 | 1 |
| *Paratrachymyrmex* nr. *bugnioni* | Par.bugn | 10 | 16 | 7 | 8 | 14 | 37 |
| *Paratrachymyrmex* nr. *diversus* | Par.dive | 1 | 0 | 1 | 0 | 1 | 0 |
| *Paratrachymyrmex* MPEG 03 | Par.s01 | 1 | 0 | 1 | 0 | 1 | 0 |
| *Paratrachymyrmex* MPEG 04 | Par.s02 | 3 | 1 | 3 | 1 | 4 | 1 |
| *Pheidole astur* Wilson, 2003 | Phe.astu | 18 | 10 | 8 | 8 | 51 | 34 |
| *Pheidole biconstricta* Mayr, 1870 | Phe.bico | 17 | 17 | 8 | 8 | 71 | 58 |
| *Pheidole bruesi* Wheeler, 1911 | Phe.brue | 0 | 3 | 0 | 5 | 0 | 10 |
| *Pheidole colobopsis* Mann, 1916 | Phe.colo | 0 | 1 | 0 | 1 | 0 | 1 |
| *Pheidole cursor* Wilson, 2003 | Phe.curs | 19 | 9 | 8 | 7 | 75 | 20 |

| | | | | | | | |
|---|---|---|---|---|---|---|---|
| *Pheidole dolon* Wilson, 2003 | Phe.dolo | 9 | 4 | 7 | 6 | 12 | 8 |
| *Pheidole fimbriata* Roger, 1863 | Phe.fimb | 2 | 2 | 2 | 1 | 2 | 2 |
| *Pheidole fowleri* Wilson, 2003 | Phe.fowl | 19 | 18 | 8 | 8 | 63 | 52 |
| *Pheidole fracticeps* Wilson, 2003 | Phe.frac | 25 | 25 | 8 | 8 | 114 | 155 |
| *Pheidole gauthieri* Forel, 1901 | Phe.gaut | 4 | 2 | 4 | 2 | 6 | 2 |
| *Pheidole jeannei* Wilson, 2003 | Phe.jean | 6 | 0 | 7 | 0 | 15 | 0 |
| *Pheidole meinerti* Forel, 1905 | Phe.mein | 18 | 10 | 8 | 6 | 31 | 16 |
| *Pheidole microps* Wilson, 2003 | Phe.micr | 1 | 1 | 1 | 1 | 1 | 1 |
| *Pheidole scolioceps* Wilson, 2003 | Phe.scol | 7 | 4 | 7 | 6 | 18 | 8 |
| *Pheidole subarmata* Mayr, 1884 | Phe.suba | 13 | 7 | 8 | 8 | 49 | 22 |
| *Pheidole synarmata* Wilson, 2003 | Phe.syna | 10 | 7 | 8 | 4 | 26 | 10 |
| *Pheidole triconstricta* Forel, 1886 | Phe.tric | 1 | 9 | 1 | 8 | 1 | 37 |
| *Pheidole vorax* (Fabricius, 1804) | Phe.vora | 2 | 1 | 2 | 1 | 2 | 1 |
| *Pheidole* nr. *araneoides* | Phe.aran | 9 | 5 | 5 | 5 | 10 | 9 |
| *Pheidole* nr. *gilva* | Phe.gilv | 0 | 1 | 0 | 1 | 0 | 1 |
| *Pheidole* nr. *pygmaea* | Phe.pygm | 22 | 17 | 8 | 8 | 80 | 51 |
| *Pheidole* (gr. diligens) MPEG 01 | Phe.s01 | 3 | 3 | 3 | 6 | 6 | 8 |
| *Pheidole* (gr. diligens) MPEG 02 | Phe.s02 | 7 | 4 | 6 | 3 | 12 | 5 |
| *Pheidole* (gr. diligens) MPEG 03 | Phe.s03 | 1 | 0 | 1 | 0 | 1 | 0 |
| *Pheidole* (gr. diligens) MPEG 04 | Phe.s04 | 1 | 0 | 1 | 0 | 1 | 0 |
| *Pheidole* (gr. diligens) MPEG 06 | Phe.s05 | 1 | 0 | 1 | 0 | 1 | 0 |
| *Pheidole* (gr. fallax) MPEG 03 | Phe.s06 | 8 | 1 | 8 | 2 | 24 | 2 |
| *Pheidole* (gr. tristis) MPEG 02 | Phe.s07 | 1 | 1 | 1 | 2 | 1 | 2 |
| *Pheidole* MPEG 01 | Phe.s08 | 7 | 2 | 6 | 2 | 12 | 2 |
| *Pheidole* MPEG 02 | Phe.s09 | 4 | 4 | 5 | 5 | 5 | 6 |
| *Pheidole* MPEG 03 | Phe.s10 | 4 | 6 | 8 | 8 | 14 | 17 |
| *Pheidole* MPEG 04 | Phe.s11 | 16 | 4 | 7 | 3 | 21 | 5 |
| *Pheidole* MPEG 05 | Phe.s12 | 0 | 1 | 0 | 1 | 0 | 1 |
| *Pheidole* MPEG 06 | Phe.s13 | 1 | 0 | 3 | 0 | 3 | 0 |
| *Pheidole* MPEG 09 | Phe.s14 | 0 | 2 | 0 | 2 | 0 | 2 |
| *Pheidole* MPEG 10 | Phe.s15 | 4 | 1 | 7 | 1 | 9 | 1 |

| | | | | | | | |
|---|---|---|---|---|---|---|---|
| *Pheidole* MPEG 11 | Phe.s16 | 4 | 1 | 5 | 1 | 10 | 1 |
| *Pheidole* MPEG 12 | Phe.s17 | 3 | 0 | 3 | 0 | 4 | 0 |
| *Pheidole* MPEG 13 | Phe.s18 | 2 | 0 | 6 | 0 | 7 | 0 |
| *Pheidole* MPEG 14 | Phe.s19 | 0 | 1 | 0 | 1 | 0 | 1 |
| *Pheidole* MPEG 15 | Phe.s20 | 1 | 0 | 1 | 0 | 1 | 0 |
| *Pheidole* MPEG 17 | Phe.s21 | 1 | 0 | 1 | 0 | 1 | 0 |
| *Pheidole* MPEG 18 | Phe.s22 | 1 | 1 | 1 | 1 | 1 | 1 |
| *Procryptocerus* MPEG 01 | Pro.s01 | 1 | 0 | 1 | 0 | 1 | 0 |
| *Rogeria lirata* Kugler, 1994 | Rog.lira | 0 | 1 | 0 | 1 | 0 | 1 |
| *Rogeria procera* Emery, 1896 | Rog.proc | 1 | 0 | 1 | 0 | 1 | 0 |
| *Rogeria scobinata* Kugler, 1994 | Rog.scob | 2 | 2 | 2 | 1 | 2 | 2 |
| *Rogeria subarmata* (Kempf, 1961) | Rog.suba | 2 | 5 | 3 | 6 | 3 | 7 |
| *Rogeria* MPEG 02 | Rog.s01 | 1 | 1 | 1 | 1 | 1 | 1 |
| *Sericomyrmex* MPEG 01 | Ser.s01 | 10 | 13 | 6 | 8 | 26 | 30 |
| *Sericomyrmex* MPEG 02 | Ser.s02 | 11 | 12 | 5 | 6 | 20 | 19 |
| *Sericomyrmex* MPEG 03 | Ser.s03 | 1 | 0 | 1 | 0 | 1 | 0 |
| *Solenopsis* nr. *iheringi* | Sol.iher | 13 | 13 | 8 | 8 | 58 | 46 |
| *Solenopsis* nr. *virulens* | Sol.viru | 5 | 0 | 6 | 0 | 12 | 0 |
| *Solenopsis* MPEG 01 | Sol.s01 | 25 | 25 | 8 | 8 | 90 | 78 |
| *Solenopsis* MPEG 02 | Sol.s02 | 20 | 20 | 7 | 8 | 38 | 34 |
| *Solenopsis* MPEG 04 | Sol.s03 | 22 | 15 | 8 | 5 | 52 | 19 |
| *Solenopsis* MPEG 06 | Sol.s04 | 2 | 1 | 4 | 1 | 4 | 1 |
| *Solenopsis* MPEG 07 | Sol.s05 | 9 | 5 | 6 | 4 | 14 | 6 |
| *Solenopsis* MPEG 08 | Sol.s06 | 2 | 0 | 1 | 0 | 2 | 0 |
| *Strumigenys carinitorax* Borgmeier, 1934 | Str.cari | 1 | 0 | 1 | 0 | 1 | 0 |
| *Strumigenys cordovensis* Mayr, 1887 | Str.cord | 0 | 1 | 0 | 1 | 0 | 1 |
| *Strumigenys denticulata* Mayr, 1887 | Str.dent | 10 | 1 | 6 | 1 | 14 | 1 |
| *Strumigenys elongata* Roger, 1863 | Str.elon | 1 | 1 | 1 | 1 | 1 | 1 |
| *Strumigenys perparva* Brown, 1958 | Str.perp | 3 | 2 | 2 | 2 | 3 | 2 |
| *Strumigenys precava* Brown, 1954 | Str.prec | 0 | 1 | 0 | 1 | 0 | 1 |

| | | | | | | | |
|---|---|---|---|---|---|---|---|
| *Strumigenys subedentata* Mayr, 1887 | Str.sube | 1 | 0 | 1 | 0 | 1 | 0 |
| *Strumigenys trudifera* Kempf & Brown, 1969 | Str.trud | 0 | 1 | 0 | 1 | 0 | 1 |
| *Strumigenys zeteki* (Brown, 1959) | Str.zete | 6 | 0 | 6 | 0 | 7 | 0 |
| *Strumigenys* (gr. schulzy) MPEG 01 | Str.schu | 0 | 1 | 0 | 1 | 0 | 1 |
| *Wasmannia auropunctata* (Roger, 1863) | Was.auro | 7 | 9 | 7 | 8 | 21 | 22 |
| *Wasmannia* nr. *lutzi* | Was.lutz | 1 | 1 | 1 | 1 | 1 | 1 |
| *Xenomyrmex* MPEG 01 | Xen.s01 | 1 | 0 | 1 | 0 | 1 | 0 |
| **Ponerinae** | | | | | | | |
| *Anochetus* MPEG 01 | Ano.s01 | 4 | 0 | 4 | 0 | 6 | 0 |
| *Anochetus* MPEG 02 | Ano.s02 | 2 | 1 | 3 | 1 | 3 | 1 |
| *Centromyrmex brachycola* (Roger, 1861) | Cen.brac | 0 | 2 | 0 | 1 | 0 | 2 |
| *Hypoponera* MPEG 01 | Hyp.s01 | 1 | 0 | 1 | 0 | 1 | 0 |
| *Leptogenys* nr. *linearis* | Lep.line | 1 | 0 | 1 | 0 | 1 | 0 |
| *Leptogenys* MPEG 02 | Lep.s01 | 1 | 1 | 1 | 1 | 1 | 1 |
| *Mayaponera arhuaca* (Forel, 1901) | May.arhu | 4 | 13 | 5 | 8 | 8 | 36 |
| *Mayaponera constricta* (Mayr, 1884) | May.cons | 11 | 22 | 8 | 8 | 27 | 74 |
| *Neoponera apicalis* (Latreille, 1802) | Neo.apic | 6 | 6 | 4 | 4 | 8 | 7 |
| *Neoponera commutata* (Roger, 1860) | Neo.comm | 14 | 5 | 8 | 5 | 22 | 5 |
| *Neoponera laevigata* (Smith, 1858) | Neo.laev | 2 | 0 | 2 | 0 | 2 | 0 |
| *Neoponera verenae* Forel, 1922 | Neo.vere | 8 | 0 | 8 | 0 | 18 | 0 |
| *Odontomachus* nr. *bauri* | Odo.baur | 11 | 0 | 6 | 0 | 17 | 0 |
| *Odontomachus* nr. *caelatus* | Odo.cael | 4 | 0 | 5 | 0 | 7 | 0 |
| *Odontomachus* nr. *meinerti* | Odo.mein | 6 | 4 | 7 | 4 | 10 | 6 |
| *Odontomachus* nr. *hematodus* | Odo.hema | 6 | 1 | 8 | 1 | 18 | 1 |
| *Odontomachus* MPEG 05 | Odo.s01 | 1 | 0 | 1 | 0 | 1 | 0 |
| *Pachycondyla crassinoda* (Latreille, 1802) | Pac.cras | 5 | 13 | 7 | 8 | 10 | 29 |

| | | | | | | | |
|---|---|---|---|---|---|---|---|
| *Pachycondyla harpax* (Fabricius, 1804) | Pac.harp | 4 | 2 | 5 | 2 | 7 | 2 |
| *Platythyrea* nr. *sinuata* | Pla.sinu | 0 | 1 | 0 | 1 | 0 | 1 |
| *Rasopone ferruginea* (Smith, 1858) | Ras.ferr | 1 | 1 | 1 | 2 | 1 | 2 |
| **Proceratiinae** | | | | | | | |
| *Discothyrea* MPEG 01 | Dis.s01 | 1 | 0 | 1 | 0 | 1 | 0 |
| **Pseudomyrmecinae** | | | | | | | |
| *Pseudomyrmex gracilis* (Fabricius, 1804) | Pse.grac | 1 | 0 | 1 | 0 | 1 | 0 |
| *Pseudomyrmex* nr. *oculatus* | Pse.ocul | 0 | 1 | 0 | 1 | 0 | 1 |
| *Pseudomyrmex* nr. *tenuis* | Pse.tenu | 1 | 3 | 1 | 4 | 2 | 5 |
| *Pseudomyrmex* MPEG 04 | Pse.s01 | 3 | 0 | 2 | 0 | 3 | 0 |
| Total register | | 902 | 702 | 619 | 492 | 2020 | 1568 |
| Total species | | | | | | 172 | 157 |

Table S3. Results of the best models (ΔAICc <2) in the temporal and spatial models for the relationship between ant species richness and covariates in the control and experimental plots. We considered the effect of each variable and double interactions. Models are ranked from best to worst according to model ΔAICc. K the numbers of estimated parameters for each model. (Litt.Div. = Litter diversity, Resp. = Respiration).

| **Scale** | **Model** | **ΔAICc** | **AICc** | **K** | **wAICc** | **$R^2$** |
|---|---|---|---|---|---|---|
| **Temporal** | Litt.Div.+ Plot + Plot:Litt.Div. | 0 | 113.08 | 6 | 0.63 | 0.66 |
| **Spatial** | Litt.Div. + Plot + Trees + Litt.Div.:Plot + Plot:Trees | 0 | 293.42 | 8 | 0.09 | 0.49 |
| | Litt.Div. + Plot + Litt.Div.:Plot | 0.37 | 293.79 | 6 | 0.07 | 0.42 |
| | Plot + Trees + Plot:Trees | 0.68 | 294.1 | 6 | 0.06 | 0.49 |
| | Litt.Div. + Plot + Trees + Litt.Div.:Plot | 0.77 | 294.19 | 7 | 0.06 | 0.46 |
| | Litt.Div. + Plot + Trees + Plot:Trees | 1.04 | 294.46 | 7 | 0.05 | 0.49 |
| | Moisture + Litt.Div. + Plot + Trees + Litt.Div.:Plot + Plot:Trees | 1.09 | 294.51 | 9 | 0.05 | 0.5 |
| | Moisture + Litt.Div. + Plot + Litt.Div.:Plot | 1.44 | 294.86 | 7 | 0.04 | 0.42 |
| | Moisture + Litt.Div. + Plot + Trees + Litt.Div.:Plot | 1.51 | 294.93 | 8 | 0.04 | 0.47 |
| | Moisture + Litt.Div. + Plot + Moisture:Plot + Litt.Div.:Plot | 1.66 | 295.08 | 8 | 0.04 | 0.43 |
| | Moisture + Plot + Trees + Plot:Trees | 1.68 | 295.1 | 7 | 0.04 | 0.5 |
| | Moisture + Litt.Div. + Plot + Trees + Moisture:Plot + Litt.Div.:Plot + Plot:Trees | 1.75 | 295.17 | 10 | 0.04 | 0.5 |
| | Moisture + Litt.Div. + Plot + Trees + Moisture:Plot + Litt.Div.:Plot | 1.91 | 295.33 | 9 | 0.03 | 0.47 |
| | Plot + Trees | 1.97 | 295.39 | 5 | 0.03 | 0.45 |
| | Litt.Div. + Plot + Trees | 1.98 | 295.4 | 6 | 0.03 | 0.46 |

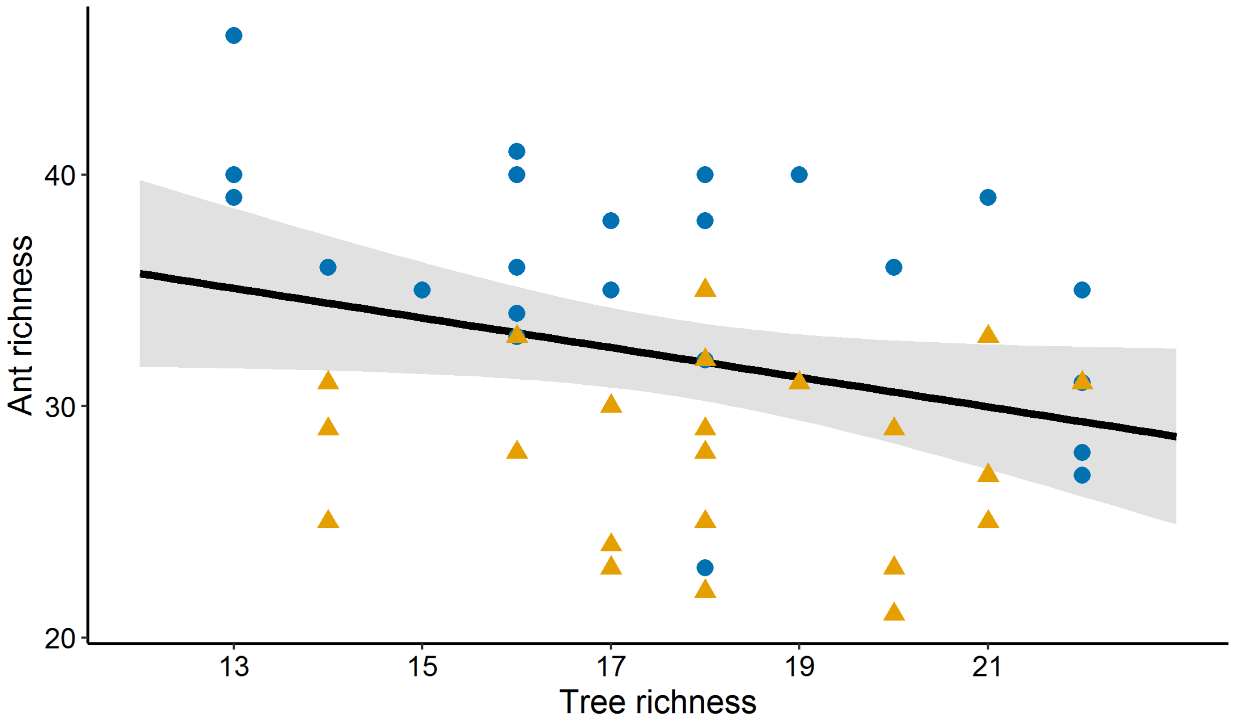


Fig. S4. Relationship between the tree and ant richness in ESECAFLOR experiment. Blue circles = control plot; yellow triangles = experimental plot.

Table S4. PERMANOVA analysis describing the relationship between ant composition and environmental variables for the temporal (moisture, litter diversity and biomass) and spatial (moisture, litter diversity, biomass, respiration, trees and DBH) models. (SS = Sum of squares; MS = Mean of squares).

| **Model** | **Predictor** | **d.f.** | **SS** | **MS** | ***Pseudo* F** | **$R^2$** | ***p**** |
|---|---|---|---|---|---|---|---|
| **Temporal** | Soil moisture | 1 | 0.35 | 0.35 | 4.85 | 0.21 | **<0.001** |
| | Litter diversity | 1 | 0.28 | 0.28 | 3.80 | 0.16 | **<0.01** |
| | Plot | 1 | 0.21 | 0.21 | 2.86 | 0.12 | **<0.01** |
| | Biomass | 1 | 0.07 | 0.07 | 0.99 | 0.04 | 0.42 |
| | Residual | 11 | 0.80 | 0.07 | | 0.47 | |
| | Total | 15 | 1.71 | | | 1 | |
| **Spatial** | Soil moisture | 1 | 0.80 | 0.80 | 3.91 | 0.07 | **<0.001** |
| | Litter diversity | 1 | 0.16 | 0.16 | 0.80 | 0.02 | 0.78 |
| | Plot | 1 | 0.38 | 0.38 | 1.86 | 0.04 | **<0.01** |
| | Biomass | 1 | 0.18 | 0.18 | 0.88 | 0.02 | 0.65 |
| | Respiration | 1 | 0.31 | 0.31 | 1.50 | 0.03 | 0.05 |
| | Trees | 1 | 0.16 | 0.16 | 0.78 | 0.01 | 0.78 |
| | DBH | 1 | 0.25 | 0.25 | 1.22 | 0.02 | 0.18 |
| | Residual | 42 | 8.61 | 0.21 | | 0.79 | |
| | Total | 49 | 10.8 | | | 1 | |

**p values* determined by permutation.

Table S5. Eigenvalues and variance explained for the first two axes in the RDA analysis for the temporal and spatial models.

| **Scale** | **Variables** | **RDA1** | **RDA2** |
|---|---|---|---|
| Temporal | Biomass | -0.21 | 0.64 |
| | Soil moisture | -0.91 | 0.03 |
| | Litter diversity | -0.15 | -0.93 |
| | Eigenvalue | 0.06 | 0.03 |
| | Explained (%) | 20.20 | 9.30 |
| | *P* | 0.001 | |
| Spatial | Biomass | -0.35 | -0.18 |
| | Soil moisture | -0.84 | 0.03 |
| | Litter diversity | -0.01 | -0.15 |
| | Respiration | 0.11 | 0.53 |
| | Trees | 0.10 | -0.34 |
| | DBH | 0.01 | -0.39 |
| | Eigenvalue | 0.03 | 0.02 |
| | Explained (%) | 6.02 | 3.70 |
| | *P* | 0.001 | |
